# Nernst Equilibrium, Rectification, and Saturation: Insights into Ion Channel Behavior

**DOI:** 10.1101/2024.08.16.608320

**Authors:** Ryan Carlsen, Hannah Weckel-Dahman, Jessica M.J. Swanson

**Author notes:** Corresponding Author **Jessica M. J. Swanson** - Department of Chemistry, University of Utah, Salt Lake City, UT, 84112 – United States.

## Abstract

The dissipation of electrochemical gradients through ion channels plays a central role in biology. Herein we use voltage responsive kinetic models of ion channels to explore how electrical and chemical potentials differentially influence ion transport properties. These models demonstrate how electrically driven flux is greater than the Nernstian equivalent chemically driven flux, yet still perfectly cancels when the two gradients oppose each other. We find that the location and relative stability of ion binding sites dictates rectification properties by shifting the location of the most voltage sensitive transitions. However, these rectification properties invert when bulk concentrations increase relative to the binding site stabilities, moving the rate limiting steps from uptake into a relatively empty channel to release from an ion-blocked full channel. Additionally, the origin of channel saturation is shown to depend on the free energy of uptake relative to bulk concentrations. Collectively these insights provide framework for interpreting and predicting how channel properties manifest in electrochemical transport behavior.

**SIGNIFICANCE:** Understanding how electrochemical potentials, such as the proton motive force, drive ion transport is a complex challenge due to the intricate coupling between electrical and chemical gradients, which can have seemingly equivalent effects. This study investigates how channel properties are differentially influenced by these driving forces to enhance or decrease directional flux. Using model systems, we demonstrate how voltage induces greater flux than the Nernstian equivalent chemical gradient yet is still perfectly balanced when the two forces oppose each other. Our findings reveal how binding site locations dictate rectification properties that flip under low and high bulk concentrations. These insights enhance our understanding of channels and transporters, providing foundational relationships between protein properties and electrochemical outcomes.

## INTRODUCTION

The protonmotive force plays a central role in bioenergetics. Like any electrochemical potential, both the chemical and electrical gradients inherent to a proton imbalance across the membrane can drive transport, and in turn many other processes, such as ATP production or drug efflux. However, understanding the relative impact of these two driving forces—one chemical and one electrical—can be challenging. It is complicated by their seemingly similar impact on charge transport and inextricable coupling that relaxes systems to their electrochemical equilibrium with either no gradients or equivalent opposing chemical and electrical forces. Herein, we explore the similarities and differences between chemically and electrically driven transport in model ion channels to help establish relationships between channel properties, such as the location and relative stability of binding sites, and the resulting electrochemical behavior, such as current, conductance, and rectification patterns.

Ion channels perform a wide variety of functions in organisms of all kingdoms of life: aiding in the propagation of nerve impulses, maintaining heart rhythm, importing metabolites, maintaining ion concentrations, and maintaining voltage gradients. (1–5) Accordingly, understanding the mechanisms by which channels open and close in response to their environment has been a central focus in the ion channel field. However, the characteristics of fully open channels are equally intriguing. For example, potassium channels like KcsA conduct potassium ions near the diffusion limit while maintaining 1000:1 selectivity for potassium ions above sodium ions, (6) despite sodium ions having a smaller ionic radius. Other channels like VDAC instead show more nuanced selectivity with a wide range of substrates. (7,8) A better understanding of open channel behavior, the focus of this work, will provide insight into these curious behaviors and add to our ability to interpret, describe, and influence channel behavior.

Conduction behavior in open channels is primarily driven by a combination of two gradients: a chemical gradient from different substrate concentrations on the two sides of an impermeable membrane and a voltage gradient consequent to the charge imbalance of all charged species. The most basic relationship between these driving forces is expressed with the Nernst potential:

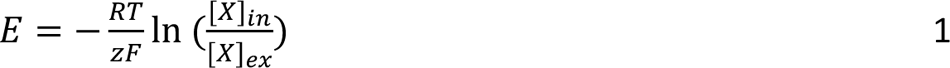

where E is the electric potential, R is the gas constant, T is temperature, z is the valence of ion X, F is Faraday’s constant, [X]_in_ is the intracellular ion concentration, and [X]_ex_ is the extracellular ion concentration. This equation is commonly used to calculate the reversal potential, since a potential of magnitude E will oppose flow for an ion concentration gradient [X]_in_/[X]_ex_. The macroscopic Poisson-Nernst-Planck theory (9) has been used for predicting ion channel currents in the last three decades (10–13) where Nernst-based continuum methods have been found to correctly predict channel currents in BD simulations driven with concentration gradients (14). While chemical gradients are relatively simple to properly incorporate into a model of an ion channel, voltage gradients have been the subject of considerable theoretical work in recent years. (15–18) Analytical methodologies have been developed that derive a voltage-coupling factor, allowing for voltage-modified free energy profiles to describe the changes in ion conduction in the open state due to an applied voltage. (19–22) The profiles can be generated for both membrane proteins and membranes themselves and clearly show unique differences from protein to protein (16,23,24). However, it is important to note that transmembrane voltage changes can also produce more significant conformational changes in a protein that would necessitate the generation of new voltage and free energy profiles. Similarly, other processes inducing major conformational changes, such as changes in pH, (25) would also require recalculation of channel profiles.

Rectification, or preferred conduction in one direction, is commonly observed in biological ion channels. (26–28) Although the most extreme examples of rectification originate from voltage gating or channel blockage, rectification is commonly observed even when measuring single channel currents in the fully open conformation. (6,29) This indicates that even when the voltage induced free energy difference in opposing directions is equal, e.g. ± 50 mV, the resulting current is not identical. Thus, the kinetics of ion transport is influenced by details beyond the net thermodynamics across the membrane.

Several factors have been discussed as originating or influencing rectification. Rectification in alamethicin channels was observed to depend on concentration and further implied to be the result of the lack of symmetry of the potential about the center (6) of the membrane. Rectification in the Shaker potassium channel was shown to change significantly with a point mutation P475D introducing a charged residue near the activation gate. (29,30) In addition, the rectification ratios of this point mutation were concentration dependent, unlike the rectification ratios in the wild-type protein. Rectification has also been observed in synthetic ion channels (31) and has been extensively explored for nanopores, where nanopore shape has been shown to induce rectification (32–35).

Herein we establish relationships between electrochemical gradients and channel properties using model systems and voltage responsive ion transport rates based on Eyring & Polanyi’s transition state theory. (36,37) This approach is not novel. Hille was the first to apply this method to sodium channels, computing IV curves with rectification. (38) Computer models based on this approach have been applied to a wide variety of channels. (30,39–42) Our focus, however, is not on reproducing experimental IV curves, but rather on understanding underlying origins of rectification, enabling experimentalists to have underlying principles to fall back on that explain why an IV curve might show rectification so that they can construct experiments to determine the specific origin in their ion channel of interest.

## METHODS

All simulated channels were constructed using a three binding site, four barrier model unless otherwise indicated. Each uptake, release, and transfer is assigned a rate based on transition state theory:

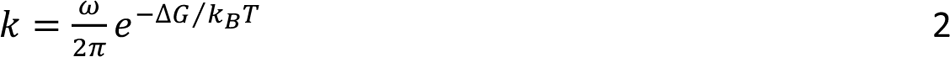

where k is the rate constant, ω is the attempt frequency (set as a constant value for all rates of 6.16975×10^10^), ΔG is the energy difference between the binding site or solution and the transition state, k_B_ is the Boltzmann constant, and T is temperature. The intrinsic rate of each transition was calculated based only on free energy (potential of mean force, PMF) energetics. Although the occupancy of neighboring binding sites will influence relative transition probabilities in real systems, these alterations were assumed to be negligible in our models to extract clear voltage-dependent relationships. The impact of concentration was folded into the PMFs by replacing the bimolecular ion uptake rates by their pseudo-first order equivalent, the product of the local concentration and the intrinsic uptake rates.

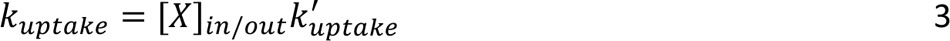

Voltage was applied as a modification to the intrinsic rate coefficients under the following assumptions. First, the transiting ions are assumed to be monovalent cations. Second, rate coefficients are adjusted for voltage following the approach of Weckel-Dahman et al. (22), based on methods established by Roux and coworkers that combine permeation free energy profiles with a dimensionless voltage coupling profile (19–21,43). For this study, we used a symmetric coupling profile φ(z) (Figure 2) consistent with a 40 Å wide membrane. The following equation describes the effect of voltage on the intrinsic rate coefficients:

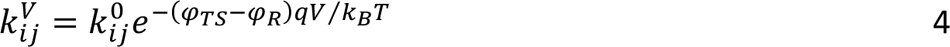

where φ_TS_ is the value of the voltage coupling factor at the transition state, φ_R_ is the value of the voltage coupling factor at the ‘reactant well’ or ion binding site, q is the ion charge, V is the voltage, k_B_ is the Boltzmann constant, and T is temperature. Note that this approach captures only the relative influence of voltage based on the locations of binding site and transition barriers. For real systems, the transition dynamics and thus rates may be additionally influenced by voltage-induced conformational changes and/or transition recrossing due to rough barriers.

**Figure 1.**
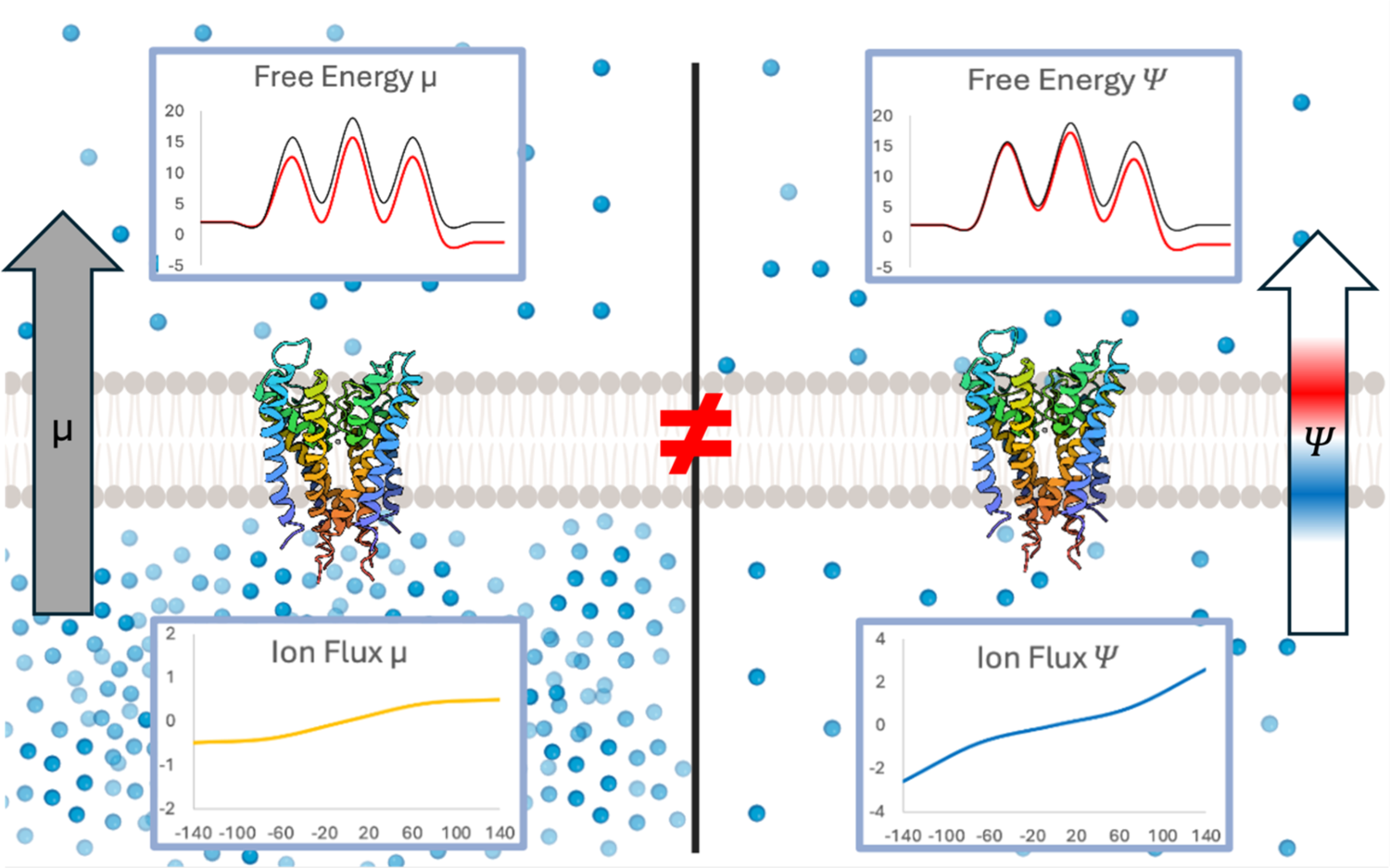
Impact of chemical gradients (left) and electric gradients (right) on free energy (top) and ion flux through the protein (bottom).

**Figure 2.**
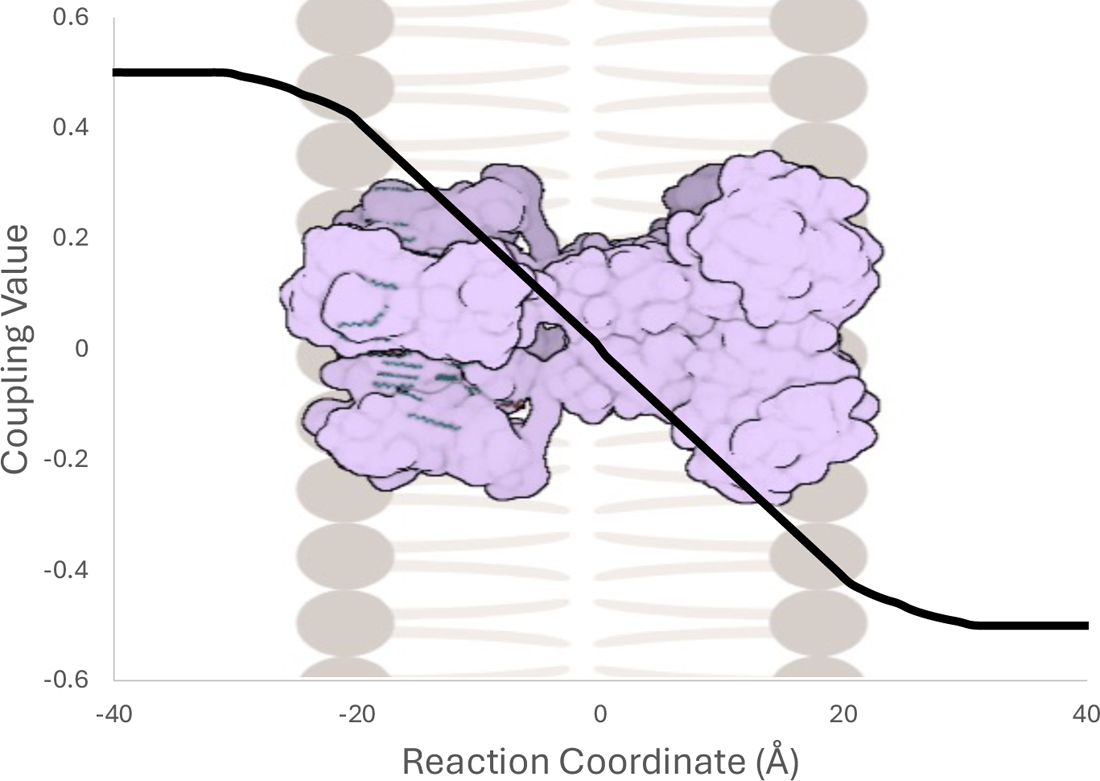
Symmetric dimensionless voltage coupling profile as a function of the membrane normal (z) reaction coordinate. Coupling factor ɸ(z) goes from a maximum value of 0.5 to a minimum value of −0.5 across a membrane width of 40 Å.

Rates, modified by voltage as appropriate, are then used to construct a rate matrix **R** for the network shown in Figure 3. Under the steady state approximation, **R** is linearly solved to determine the populations of each system state. State populations are then used to determine the system flux f, as described in further detail in the methods by Weckel-Dahman et. al. (22) Flux is signed such that positive ions flowing from the intracellular solution to the extracellular solution is positive.

**Figure 3.**
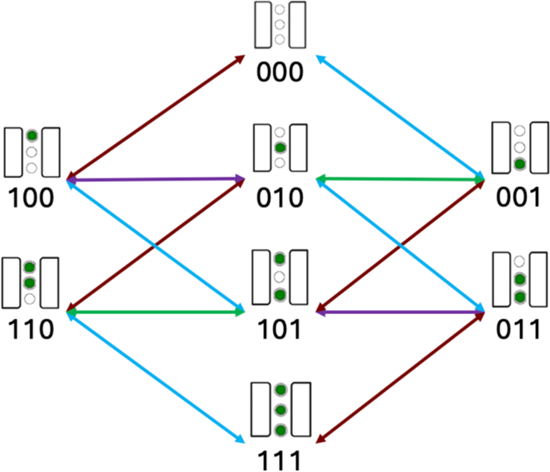
Kinetic network used to model flux through ion channels. All edges of the same color represent rates with the same magnitude in the symmetric model. Forward and reverse rates have unique values. Ion binding in the channel is represented schematically above the binary string, with the upper site and left-most digit being representing the intracellular side, respectively.

Rectification ratios (RR) are used to describe the degree to which flow is rectifying (preferred in a single direction at an absolute voltage magnitude) independent of the magnitude of the flux. The following equation was used to calculate RR:

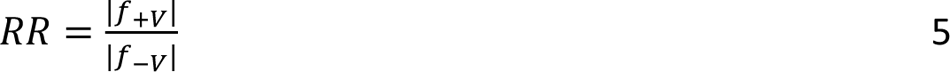

where the +/- symbols indicate the sign of same voltage magnitude. RR > 1 corresponds to greater flow outward, or outward rectification, and RR < 1 corresponds to inward rectification.

## RESULTS AND DISCUSSION

### Chemical vs. Voltage Transformation

The movement of charged substrates through channels and transporters typically involves multiple rare event transitions separating metastable intermediates. The frequency of these transitions depends on the transition ensemble dynamics and is nontrivial to fully characterize but can be estimated based on transition state theory from the free energy profiles separating metastable intermediates. A potential of mean force (PMF) curve condenses the complex transition free energy landscape into an easily interpretable one- or two-dimensional form by integrating out the remaining dimensions. For the purpose of unraveling fundamental trends electrochemically driven transport, we constructed a model PMF (Figure 4A) to describe a simple three-binding site ion channel. This PMF is symmetrical along the membrane normal and centered at the midpoint of the membrane. Each binding site has identical binding free energy, and the transition barriers are of equal height relative to the binding sites. Only the intracellular binding and extracellular binding transitions have unique barriers.

**Figure 4.**
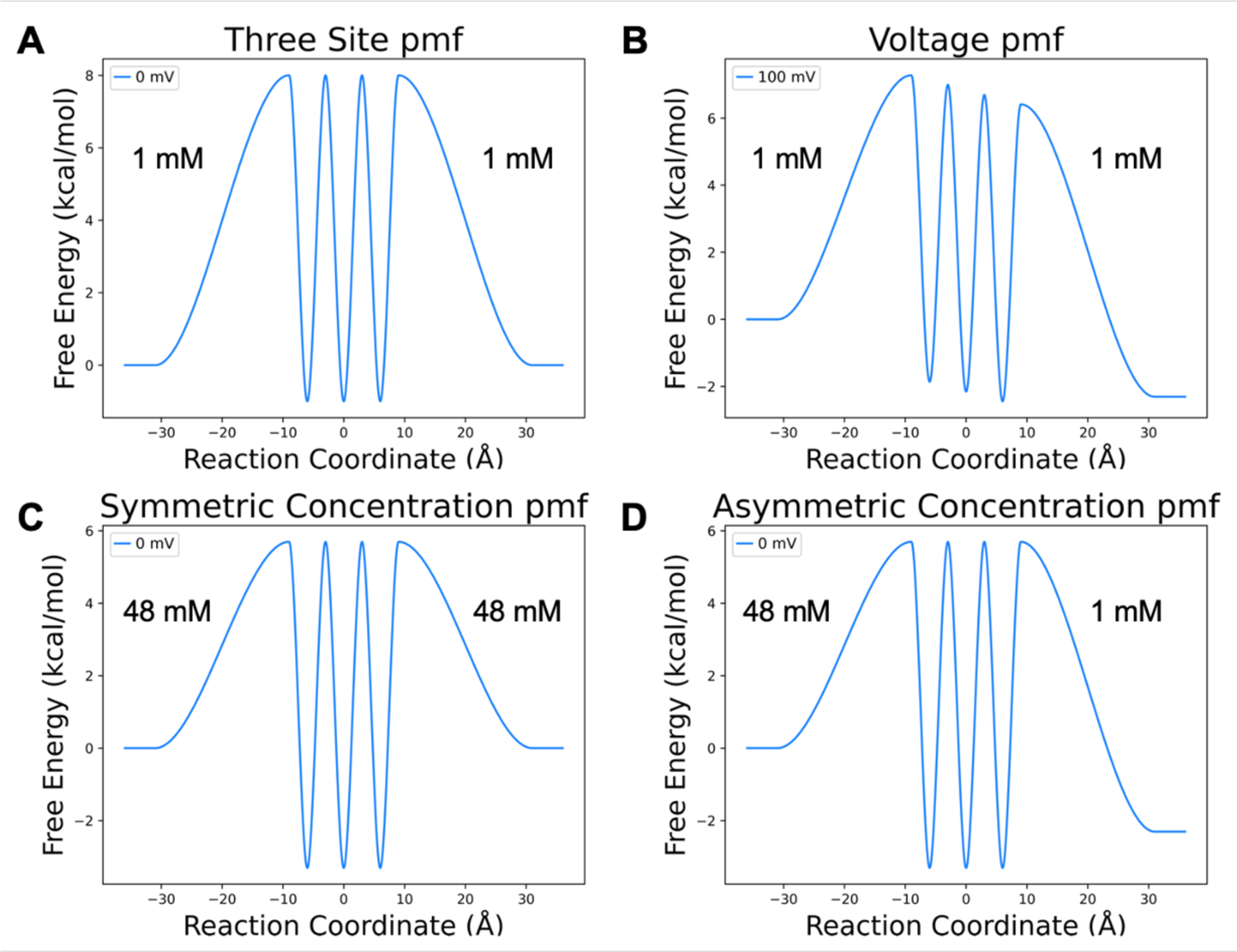
Model PMFs for an ion channel with three binding sites. A) Baseline symmetric PMF with 8 kcal/mol barriers, −1 kcal/mol binding site stabilities, and symmetric 1 mM intra/extracellular concentrations. B) Baseline PMF transformed by an applied voltage C) Baseline PMF transformed by symmetric increase in concentration. D) Baseline PMF transformed by asymmetric concentration. The 48:1 mM concentration ratio is selected to give an identical free energy gap to 100 mV applied voltage.

It is first important to recognize the differences between the two key driving forces—voltage and substrate concentration. While the Nernst relationship asserts that flux through a channel driven by a chemical gradient can be counteracted by the appropriate opposing electrical gradient, the transformations on the PMF by these two influences are not the same. In the case of voltage, (Figure 4B) the free energy profile is shifted in roughly linear, though system-specific, (19,20) manner, altering both the relative stabilities of binding sites and rates of transitions along the channel pore. Over the distance between each local minimum and the transition state, the ion will either gain or lose relative probability (free energy) through interaction with the electric field, which effectively raises or lowers the transition barrier. Thus, the electric field alters the rates of each transition that involves displacement along the membrane normal through an exponential relationship Eq. 4. In contrast, concentration gradients only change the relative population of ions on either side of the membrane Eq. 3. Thus, concentration gradients influence uptake rates through a linear relationship. They do not, however, alter the relative stability of intermediate binding sites, nor transition states other than uptake.

Consistent with these principles, our PMFs are constructed with the ends referencing the relative concentrations (e.g., 100 mM:10 mM) instead of the standard state (1M). It follows that a change in the local concentration either stabilizes or destabilizes the free energy of the bulk. For symmetric concentrations, this effectively raises or lowers the energy of the three binding sites relative to the bulk (Figure 4C), with increasing symmetric concentration effectively stabilizing binding sites inside the channel and v.v.. For asymmetric concentrations, only one uptake rate in the PMF is shifted, leaving the shape of the remainder of the PMF unaltered (Figure 4D).

### Ion flux under voltage and chemical gradients

To observe how binding site stabilities and locations impact substrate flux through the channel in combination with chemical and voltage gradients, we use a simple two binding site channel that transports a monovalent cation through a membrane (Figure 5A). This immediately captures how the ion fluxes being driven by the electric and chemical driving forces are not equivalent, even when the bulk-to-bulk free energy perturbation from the forces have equivalent Nernstian magnitudes (Figure 5B & 5D). As the electric force is increased, the flux through the protein increases in a non-linear manner producing a hyperbolic sine curve that steeply increases with increasing voltages (Figure 5B Blue line). In contrast, the chemical driving force is a hyperbolic tangent, with the slope of the line decreasing with increasing concentration gradients (Figure 5B Yellow). This is a direct consequence of flux being directly related to concentrations and exponentially related to voltage-induced rate changes. Despite this apparent disparity in the magnitude of flux for the two driving forces, they appropriately cancel out to abolish net flux once they are set in opposing directions (Figure 5B Green line). Two Nernst equivalence points demonstrate this cancellation of flux (what would be observed at +/= reversal potentials) in free energy profiles with no free energy gap between intracellular and extracellular solutions. However, the binding site stabilities are indeed shifted, (Figure 5D) leading to different equilibrium populations (Figure 5C, SI Tables S5 – S8).

**Figure 5.**
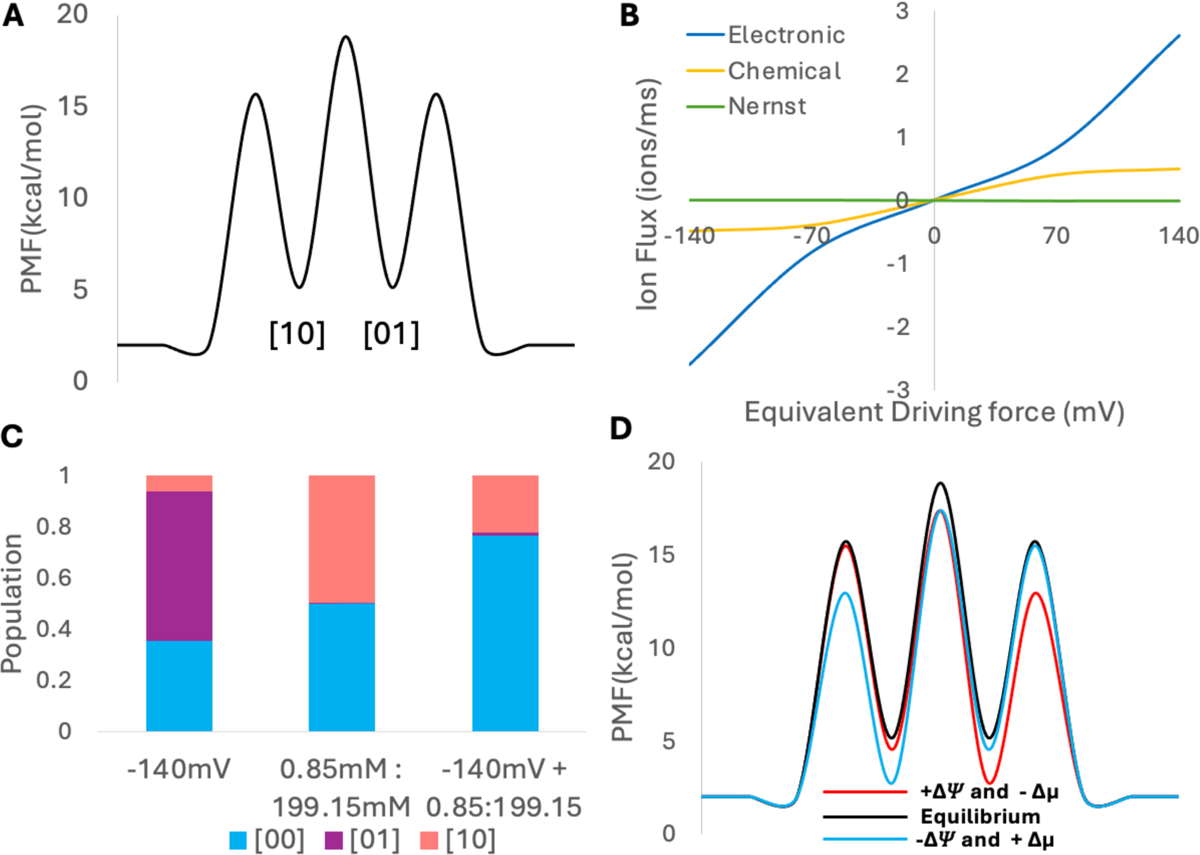
A) PMF of two-site symmetric system with symmetric 100 mM concentrations. B) Ionic flux through channel under different conditions: +140 mV electronic potential (blue), the Nernstian equivalent chemical concentration gradient [199.15 mM] ext: [0.85 mM] int (yellow), and a +140 mV electronic potential combined with a [0.85 mM] ext: [199.15 mM] int chemical gradient (Nernst equilibrium, green). C) Populations for conditions in (B). D) PMF of Nernst equilibrium at −140 mV (blue) and +140 mV (red), highlighting unique equilibrium populations and barrier heights.

The locations of the binding sites and transition states along the membrane normal controls the magnitude by which an applied transmembrane voltage speeds up or slows down transition rates. (22) To test if the chemical vs voltage flux disparity is consistent as site and barrier locations change, we constructed three additional two-site models: an asymmetric binding site model (See SI Figure S1 A, SI Tables S2 and S6), a symmetric binding site model with more stable binding sites than the bulk (See SI Figure S1 B, SI Tables S3 and S7), and an asymmetric model with less stable binding sites than the bulk (See SI Figure S1 C, SI Tables S4 and S8). Additionally, we moved the site locations for the symmetric binding site model with more stable sites by 10 angstroms to observe if this change would affect the large flux discrepancy between chemical and electric driving forces (SI Tables S9 to S12). The electric driving force consistently moved ions through the transporter with a greater flux than the chemical counterpart, but ultimately canceled out when opposing forces were combined via the Nernst (See SI Tables S9 to S12).

Despite different magnitudes of flux between chemical and electric driven processes, the Nernst-Planck theory holds under all models tested. This tells us that the interplay between ion gradients, location of the binding sites and site stability gives each protein a unique Nernstian response, with a unique free energy profile and populations.

### Site Location

Our observations of Shaker and related potassium channels (22) indicated that the site location of the binding sites and transition states would also influence the rectification behavior of ion channels. Shaker notably has its selectivity filter, containing at least three critical ion binding sites for potassium selectivity, located very near the extracellular membrane. Shaker is also inwardly rectifying, a common trait for potassium channels. We reasoned that this clustered location of binding sites may contribute to the rectifying behavior.

First, starting from a symmetric PMF, we tested additional symmetric combinations, changing the spread of the binding sites, the binding site stabilities, and the transition barrier heights. None of these symmetric operations on the PMF introduced any degree of rectification, with rectification measured using the RR (Eq. 5). (SI Figure S2 and S3)

Having established that symmetry could not produce rectification, we introduced asymmetry by pulling the entire set of sites toward intracellular side, introducing an overall 5 Å displacement in .25 Å intervals. Transition state barrier locations were maintained at the halfway point between site and membrane (± 20 Å) or site to site. This transformation was applied both with binding sites that were stable relative to the bulk concentrations, mimicking high bulk concentrations, and with similarly unstable binding sites, mimicking low bulk concentrations (Figure 6A & 6B).

**Figure 6.**
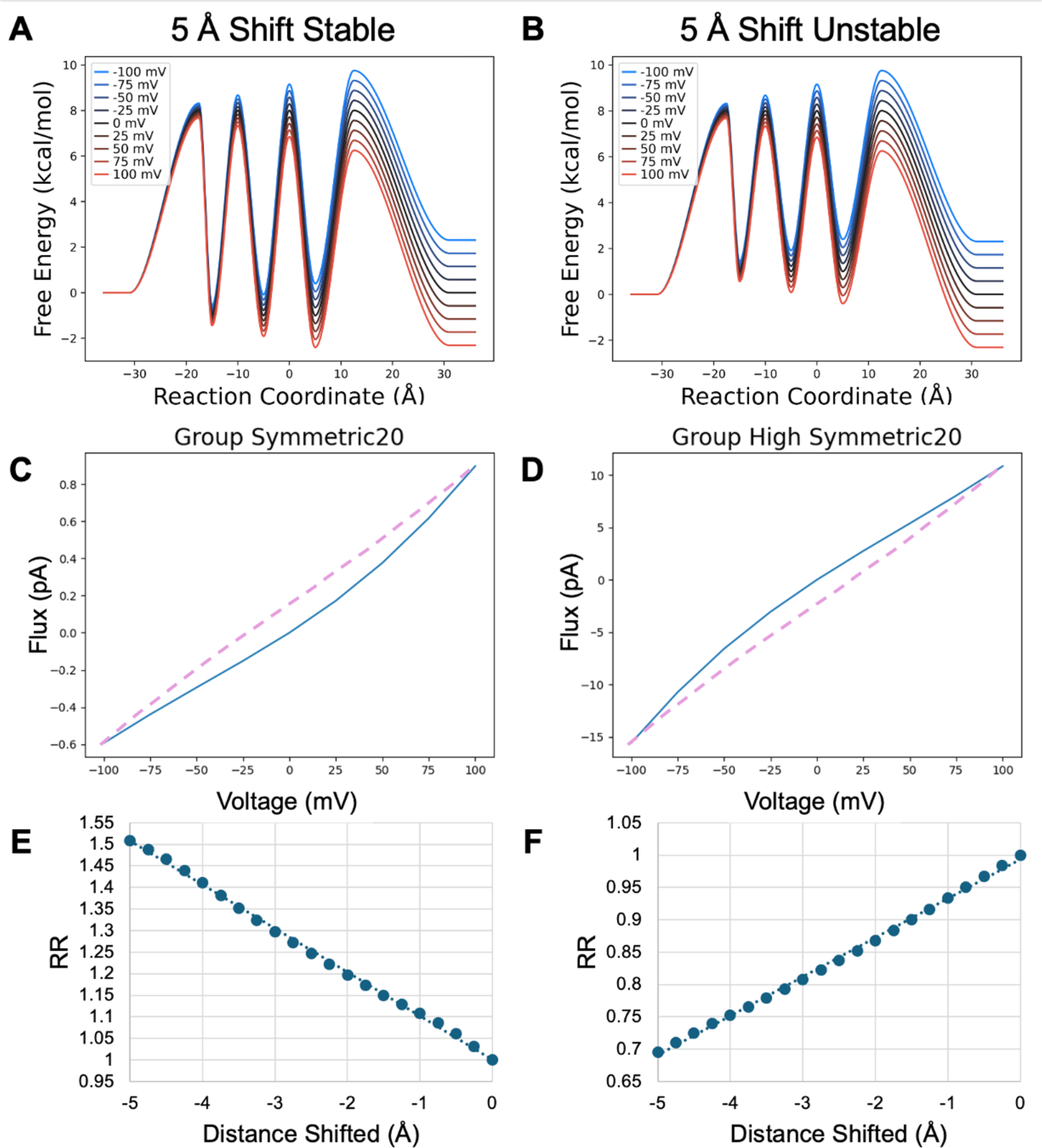
Model PMF transformed by shifting all binding sites toward the intracellular membrane. Barriers are positioned midway between binding sites or midway between a binding site and the membrane surface. A, B) Fully shifted PMFs with stable and unstable binding sites C, D) IV curves for the fully shifted PMFs. Purple dashed curve is from the untransformed PMF for comparison. E, F) Rectification ratios as the binding site locations are shifted by 0.25 Å increments to the 5 Å maximum shift.

Importantly, the relative stability of the binding sites reverses the direction of rectification (Figure 6C & 6D), illuminating why concentration-dependent rectification is commonly observed. This results from the fact that altering the stability of the binding sites changes the location of the rate limiting step.

When the binding sites are more stable than bulk, each transfer is the rate limiting step, such that there is a buildup of ions in the channel. Thus, the flux is aided primarily by speeding up release and slowing down uptake on the side toward which flux occurs. When the sites are moved to the inside, the final release, from the third site to extracellular bulk, and uptake from extracellular bulk are most influenced by voltage since they involve a greater distance between their reactant minima and transition state. Thus, they have the largest shift in activation energy. This removes the last ion blocking transport with increasing positive voltage and is the cause of outward rectification.

Conversely, when the binding sites are unstable relative to bulk, uptake is rate limiting. Thus, moving the binding sites toward the intracellular side encourages extracellular uptake while similarly decreasing intracellular uptake. This produces inward rectification. In both cases the RR changes in an approximately linear fashion with respect to site location, which may be useful for testing these models against the behavior of potassium channels, which show both differences in the location of their selectivity filter along the membrane normal and the degree of rectification observed.

To further test this initially observed trend, asymmetry in site location was also introduced by moving only the first barrier toward the intracellular side and by moving both the first barrier and the first binding site toward the intracellular side (Figure 7). When the binding sites are more stable than bulk moving the first barrier alone produces inward rectification by increasing the speed of the rate limiting release under negative voltages (unblocking the ion traffic jam). However, the magnitude of this effect is diminished compared to moving all sites and transitions since moving the first barrier to the left also decreases the voltage sensitivity of uptake from intracellular bulk, thereby increasing the speed of the reverse step that counteracts inward flux. In contrast, moving the first barrier and binding site to the left results in outward rectification, since the voltage sensitivity of release to the outside is now greater than that to the inside and the counteracting uptake from the inside under negative voltages decreases inward flux.

**Figure 7.**
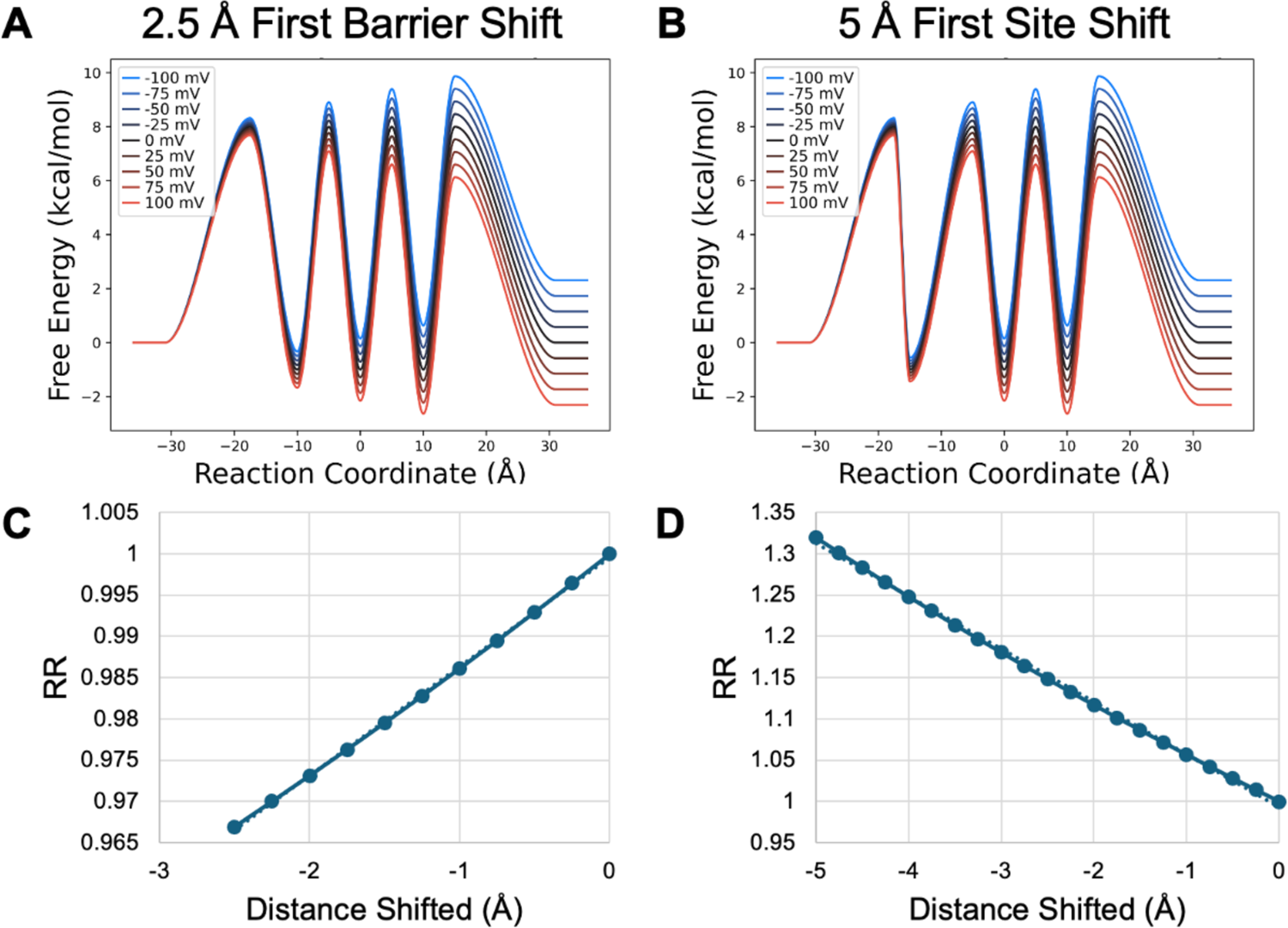
Model PMF with stable binding sites transformed by shifting A) the first barrier toward the intracellular membrane by 2.5 Å and B) the first binding site toward the intracellular membrane by 5 Å, with the first barrier also shifting to remain at the midpoint between the first binding site and the membrane surface. C, D) RR as a function of shifting distance in 0.25 Å increments.

When binding sites are unstable and uptake is rate limiting, slowing uptake from the inside becomes the dominant effect—diminishing the extent to which positive voltage can push outward flux, resulting in inward rectification whether only the barrier is moved or the barrier and binding site are moved. The degree of inward rectification is greater when both barrier and binding site are moved because the outward flux opposing release to the inside also has decreased voltage sensitivity and is thus relatively faster than if only the barrier is moved.

Both of these examples demonstrate the same principles seen for moving the entire group of barriers and binding sites—rectification results from altered voltage sensitivity on the rate limiting steps. When binding sites are more stable (high bulk concentrations), an ion blockade is established and release is rate limiting. When sites are less stable (low bulk concentrations), uptake is rate limiting. In either case, increasing voltage sensitivity by increasing the distance between the rate-limiting minimum and transition barrier increases flux leading to increased current with either positive voltages (outward rectification for a cation) or negative voltages (inward rectification).

### Peak Height and Rectification

Having explored the effects of site location, we next wanted to understand how asymmetric energetics of the PMF might influence rectification. First, we tested how the height of the initial barrier influences rectification, with both stable and unstable binding sites. With stable binding sites (Figure 8), when release is generally rate limiting, we observe only modest inward rectification when the first barrier is reduced (Figure 8 A,B). This is predominantly due to decreasing the barrier of the last release step for inward flux from 8 kcal/mol gradually down to 5 kcal/mol. This effect quickly levels off because the other transfer steps are still rate limiting. Although the RR’s look small, they are on par with the values demonstrated above. However, when the first barrier is raised, there is a much more dramatic increase in the RR’s due to outward rectification. This is again due to the last release step toward the intracellular side, but in this case, influx is being rapidly shut down as this barrier grows larger than all other barriers in the channel.

**Figure 8.**
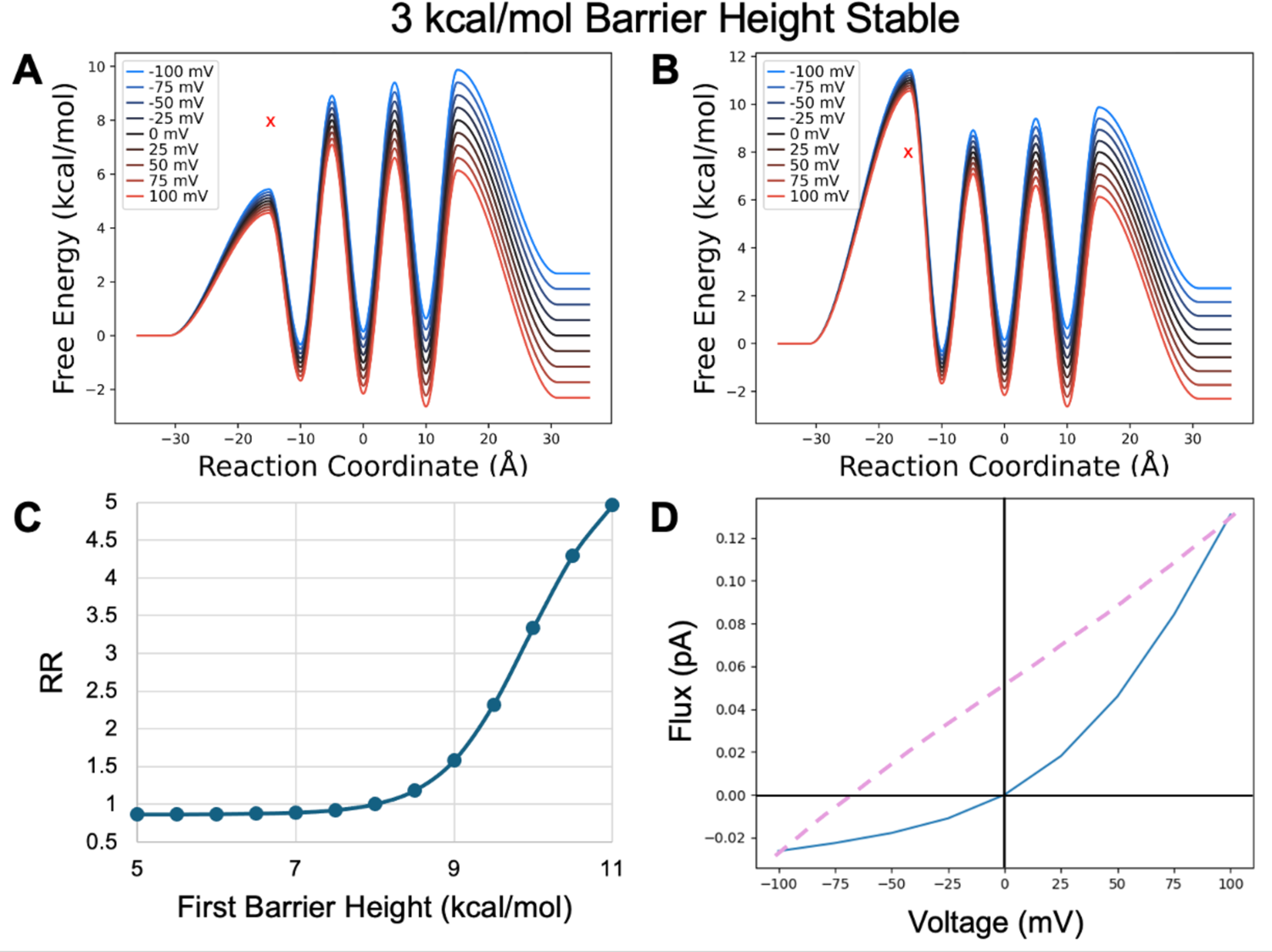
Model PMF with stable binding sites transformed by changing the first barrier height from A) a minimum value of 5 kcal/mol to B) a maximum value of 11 kcal/mol in 0.5 kcal/mol increments. X marks original barrier height. C) RR as a function of barrier height transformation. D) IV curve for the PMF shown in B.

We see this same type of behavior when we alter the height of the initial barrier with unstable binding sites (Figure 9). Here we clearly note the sigmoidal shape of the RR curve relative to the height of the initial barrier, showing an asymptotic leveling off as the other transitions limit flux. We also note that since the rate limiting step is now uptake, the direction of the rectification is mirrored compared to our PMF with unstable binding sites when release was limiting. As you decrease the rate limiting uptake barrier, you enable increased outflux (outward rectification) until the other transitions limit this increase. As you increase the rate limiting barrier, we see decreased outflux relative to influx (inward rectifying) again until the other transitions dominate.

**Figure 9.**
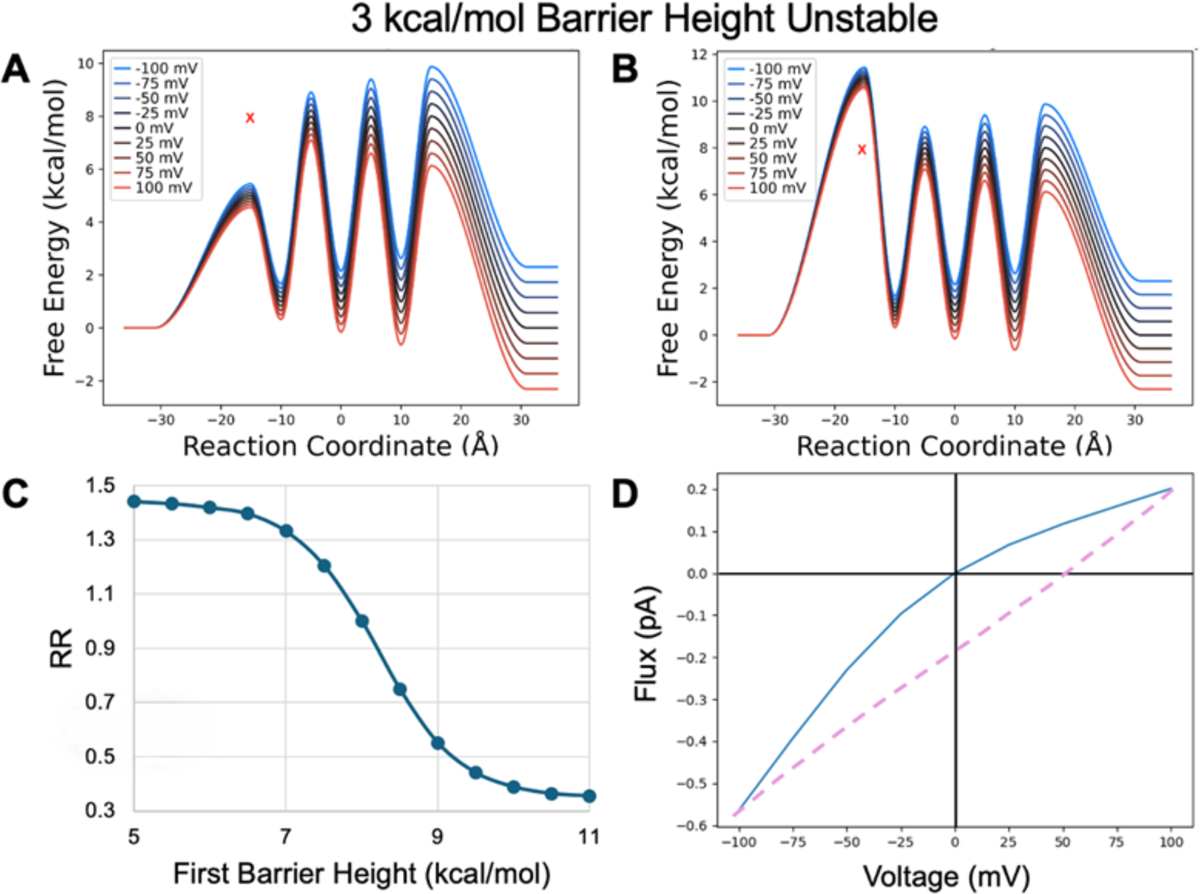
Model PMF with unstable binding sites transformed by shifting the first barrier height from A) a minimum value of 5 kcal/mol to B) a maximum value of 11 kcal/mol in 0.5 kcal/mol increments. X marks original barrier height. C) RR as a function of barrier height transformation. D) IV curve for the PMF shown in B.

When the first binding site has its depth altered, the same underlying trends govern the observed rectification (Figure 10). When the sites are more stable than bulk, increasing the stability of the first binding site dramatically slows down inward flux, rapidly increasing the RR and causing outward rectification. This is consistent with the reverse step (that opposing flux) being slower for outflux than influx (the barrier from site two to site 1 is larger than from intracellular to site one). Again, the effect is more dramatic because this creates the largest barriers in the system. In contrast decreasing its stability speeds release to the inside, leading to modest inward rectification that quickly levels off as the other rate limiting transitions dominate.

**Figure 10.**
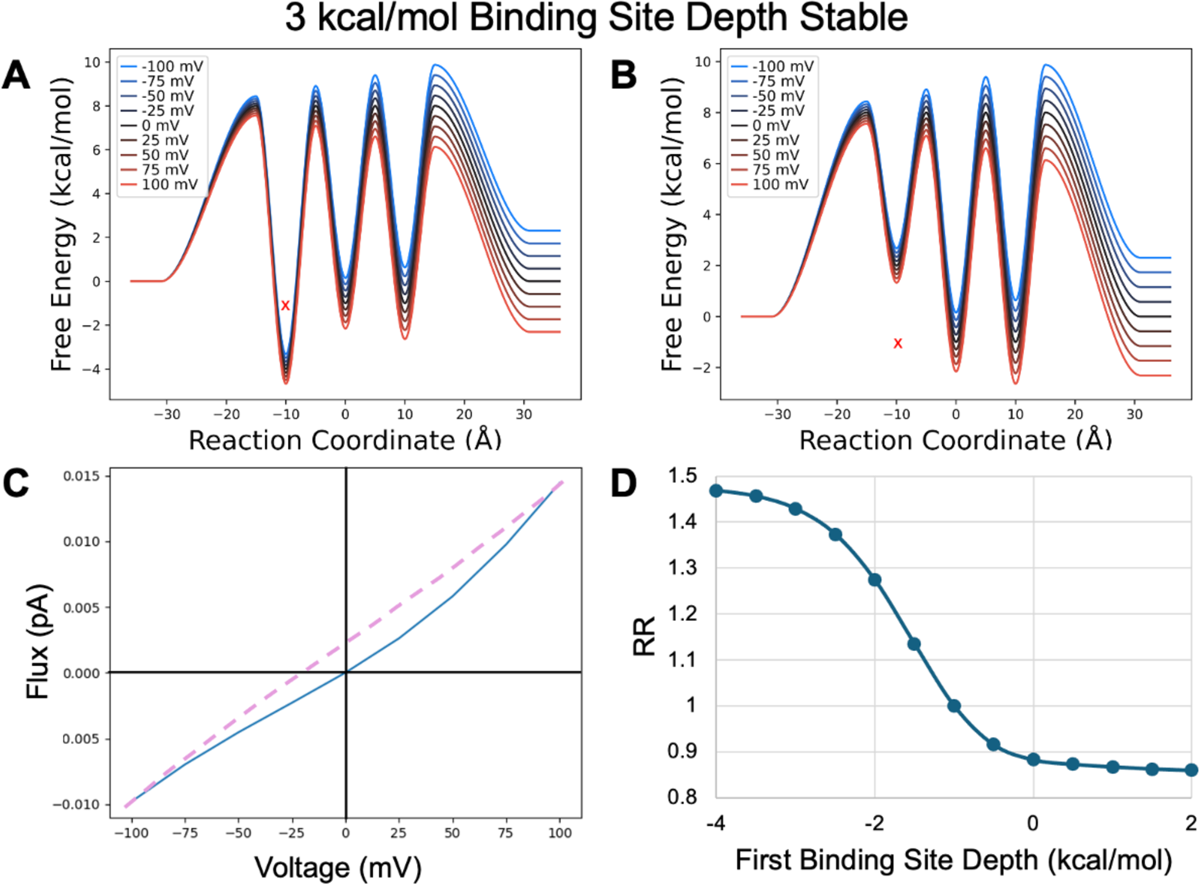
Model PMF with stable binding sites transformed by shifting the first binding site depth from A) a minimum value of − 4 kcal/mol to B) a maximum value of 1 kcal/mol in 0.5 kcal/mol increments. X indicates original binding site depth C) IV curve for the PMF shown in A. D) RR as a function of binding site depth transformation.

When the binding sites are unstable relative to bulk (high bulk concentrations, Figure 11) the rectification behavior is again flipped compared to above (low bulk concentrations). When the first site is stabilized it again becomes rate limiting in both directions. However, now the slower reverse step is for influx (intracellular to site one) leading to inward rectification. When the first site is destabilized, the second transition speeds up and the reverse step is slower for outflux, causing outward rectification that is quickly tempered by the other transition barriers.

**Figure 11.**
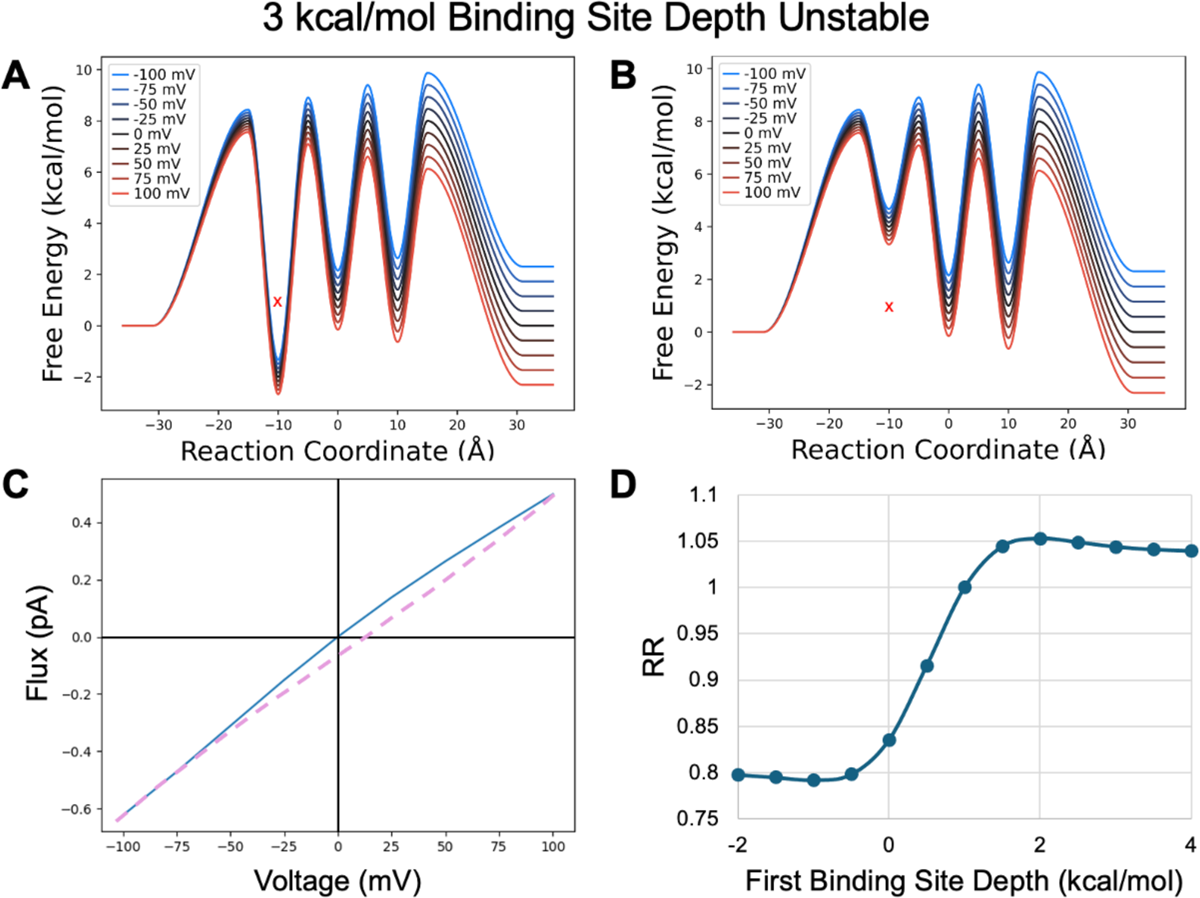
Model PMF with unstable binding sites transformed by shifting the first binding site depth from A) a minimum value of −2 kcal/mol to B) a maximum value of 4 kcal/mol in .5 kcal/mol increments. X indicates original binding site depth. C) IV curve for the PMF shown in A. D) RR as a function of binding site depth transformation.

### Saturation behavior in ion channels

It has been demonstrated for many ion channels that increasing flux through the channel as symmetric concentration is increased eventually stops (6,29) a phenomenon known as saturation. For some channels, it has even been observed that increasing symmetric concentrations of ions can lead to decreases in flux through the channel. (44) As seen in Figure 4C, this may be the result of binding sites becoming too stable relative to the intracellular and extracellular solutions, resulting in a decrease in flux as the rate of counter-flux uptake becomes much greater than the rate of transfer, blocking ion release and productive flux.

To further explore this trend, we generated Iμ curves as an analogue to IV curves so that we could clearly delineate how concentration affects flux across a wide range of values (Figure 12). Starting from a symmetric PMF with unstable binding sites (i.e., the low concentration limit), we can see that the flux through the channel first increases, reaching a maximum at 39 mM, and then slowly decreases from that point on. From the relationship between free energy and concentration, we can calculate that the binding sites’ transition from being unstable relative to the solution at 29 mM, which we initially predicted would be the point at which flux would reach its maximum, however we clearly observe that increasing concentration continues to increase flux beyond that point.

**Figure 12.**
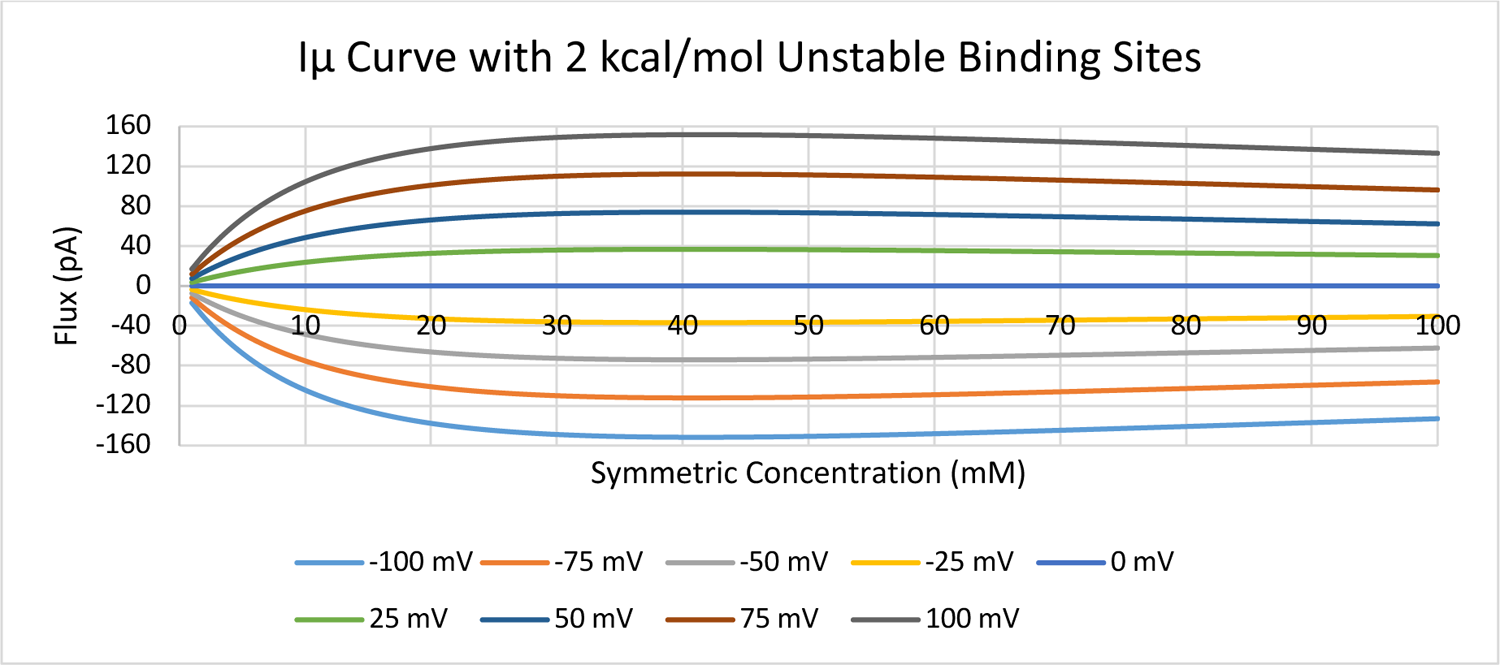
Current-chemical potential (Iμ) curve for symmetric concentration increases in 1 mM increments from 1 mM to 100 mM across a range of voltages. Underlying PMF is equivalent to Figure 4A except with binding site stabilities set at 2 kcal/mol.

The turnaround is even more pronounced (Figure 13) as the binding sites are further stabilized, only unstable relative to the solution by 1 kcal/mol. However, it is important to note that by changing the stability of the binding sites, we have also increased the height of the transfer barriers, keeping their relative height to solution. Again, we observe that despite the binding sites becoming more stable than solution at 5 mM, the maximum rate is at 7 mM. Additionally, the curvature of the individual corresponding IV curves for each concentration changes (Figure 14), beginning with super-ohmic curvature, then transitioning to being ohmic, and slightly sub-ohmic, and then back through ohmic to super-ohmic again, all as a function of the changing concentration. Interestingly, once binding sites become more stable than the bulk ionic concentrations, flux immediately decreases with increasing concentration (SI Figure S4). However, since the decrease is logarithmic, it is likely often missed in experimental assays or assumed to be within measurement noise. Collectively, the subtle interplay between how the concentration affects the populations of the binding sites and the voltage changing the heights of the barriers provides further insight into mechanisms nature can employ in designing the exquisite details of ion channels.

**Figure 13.**
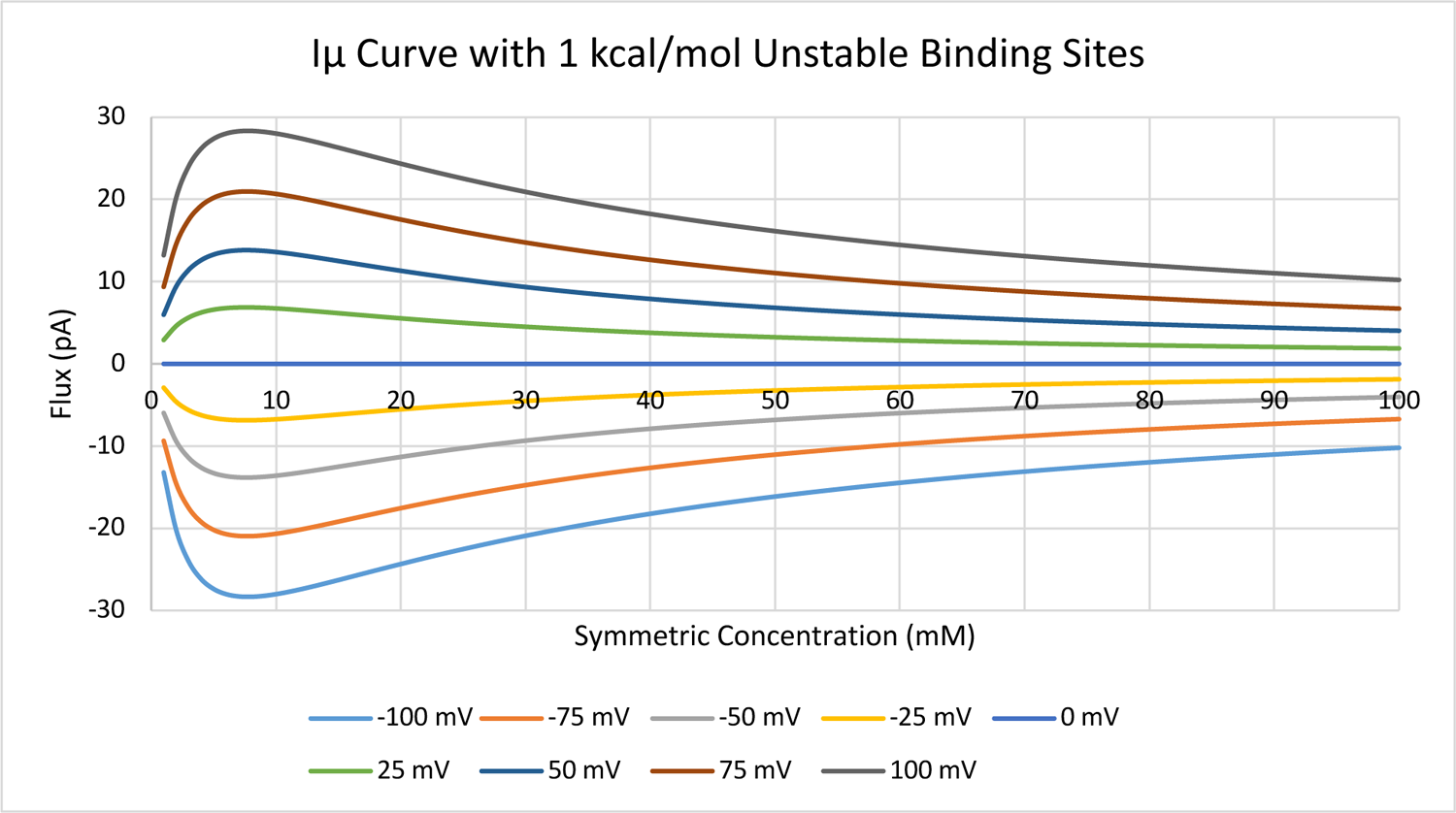
Current-chemical potential (Iμ) curve for symmetric concentration increases in 1 mM increments from 1 mM to 100 mM across a range of voltages. Underlying PMF is equivalent to Figure 4A except with binding site stabilities set at 1 kcal/mol.

**Figure 14.**
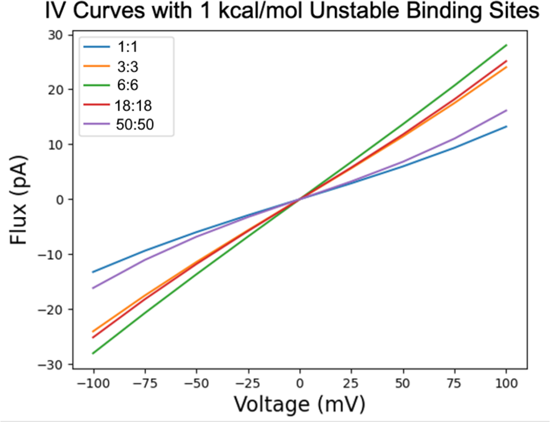
IV curves at selected concentrations for the same PMF modeled in Figure 13. Concentration curves show a behavior transformation from superohmic at the lower concentration, toward ohmic, then becoming subohmic before returning back to superohmic behavior at high symmetric concentrations.

## CONCLUSION

This work has explored the fundamental relationships between electrochemical gradients and channel properties, revealing trends that apply broadly to any transmembrane system with spatially resolved binding sites for charged substrates. The presented model systems capture how chemical and voltage effects on channels are unique, altering the PMF in substantively different ways. This is particularly obvious when the Nernst reversal potential relationship between a chemical gradient and the corresponding reversal voltage is investigated in detail, revealing that while equivalent free energies must oppose each other when at odds, it does not follow that they produce equivalent flux when separated.

Rectification is shown to arise from asymmetry in the PMF, which can be induced in a variety of ways. The relative stability of the binding sites and bulk are essential to understanding the direction of rectification. As a rule of thumb, when binding sites are more stable than intracellular and extracellular solutions (high bulk concentrations), ion occupancy in the channel grows, and the release steps become rate limiting. This is analogous to a traffic jammed freeway, wherein ions waiting to transition to the next binding site are blocked by those ahead. In contrast, when binding sites are less stable (low bulk concentrations), uptake becomes rate limiting. This could be related to a relatively empty freeway with overzealous traffic control lights on freeway entrances. Increasing the voltage sensitivity of these rate limiting steps then leads to faster forward flux and slower back flux as a function of voltage, determining the direction and magnitude of rectification. In this manner, changing the relative stability of the channel binding sites with respect to intracellular and extracellular concentrations inverts the direction of rectification by changing the location of the rate limiting steps. Concentration-dependent rectification is natural consequence of these relationships and can be used to better interpret the underlying causes of rectification for a given system of interest.

Saturation behavior is also explored and again demonstrated to be related to the stability of binding sites relative to intracellular and extracellular solutions. Generally, flux will increase with increasing concentrations until and slightly beyond when the binding sites become more stable than bulk concentrations, and thereafter will slowly/logarithmically decrease. Future work will explore how including ion-ion electrostatic interactions and limiting network connectivity, and hence multi-pathway flux, alters the relationships observed herein.

## AUTHOR CONTRIBUTIONS

R.C, H.W.D, and J.M.J.S designed the research. R.C. and H.W.D. built and analyzed the kinetic models. All authors helped interpret the results. J.M.J.S directed and funded the research. All authors wrote the manuscript and agreed on the final version.

## ACKNOWLEDGMENT

This work was supported by NIH NIGMS (R35GM143117) and the computational resources provided by the Center for High-Performance Computing (CHPC) at the University of Utah.

## TOC

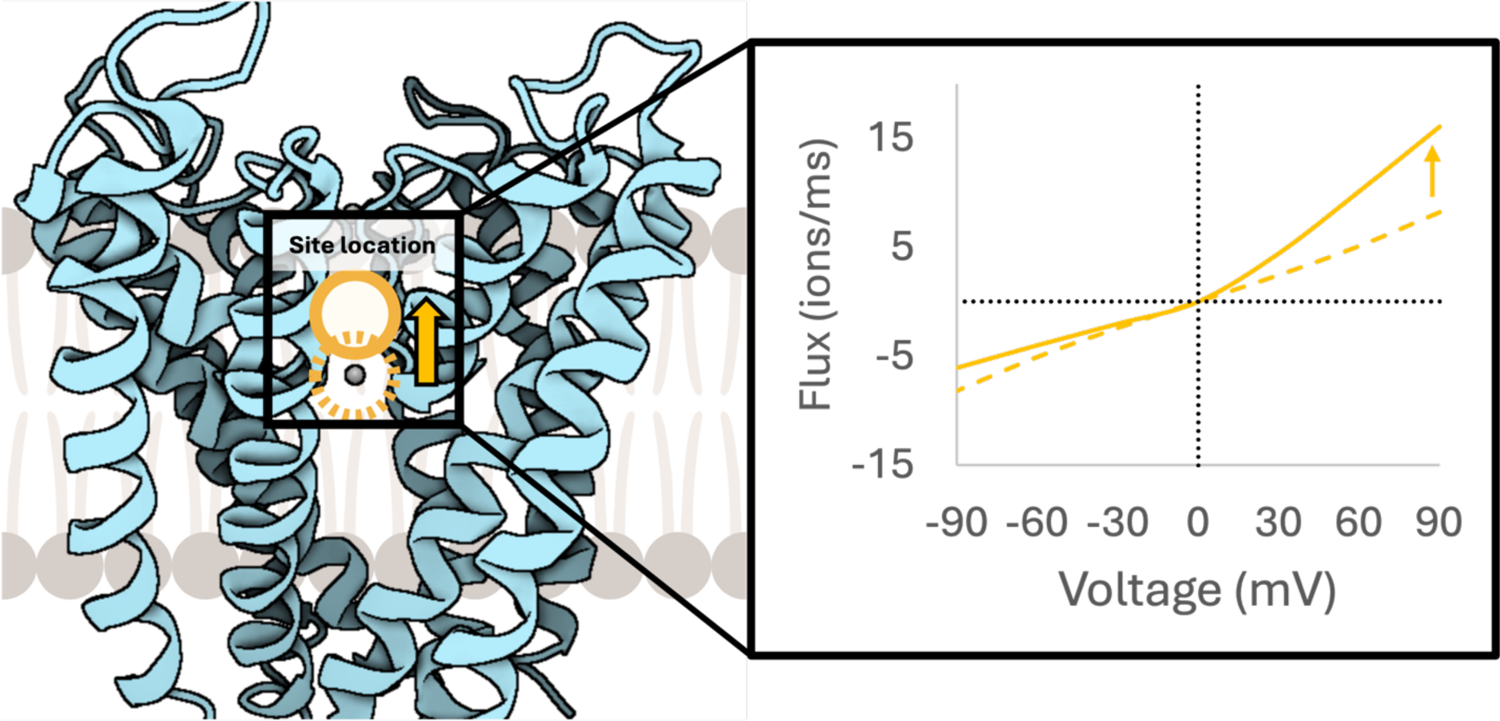

