## Supplementary Information for "Nernst Equilibrium, Rectification, and Saturation: Insights into Ion Channel Behavior"

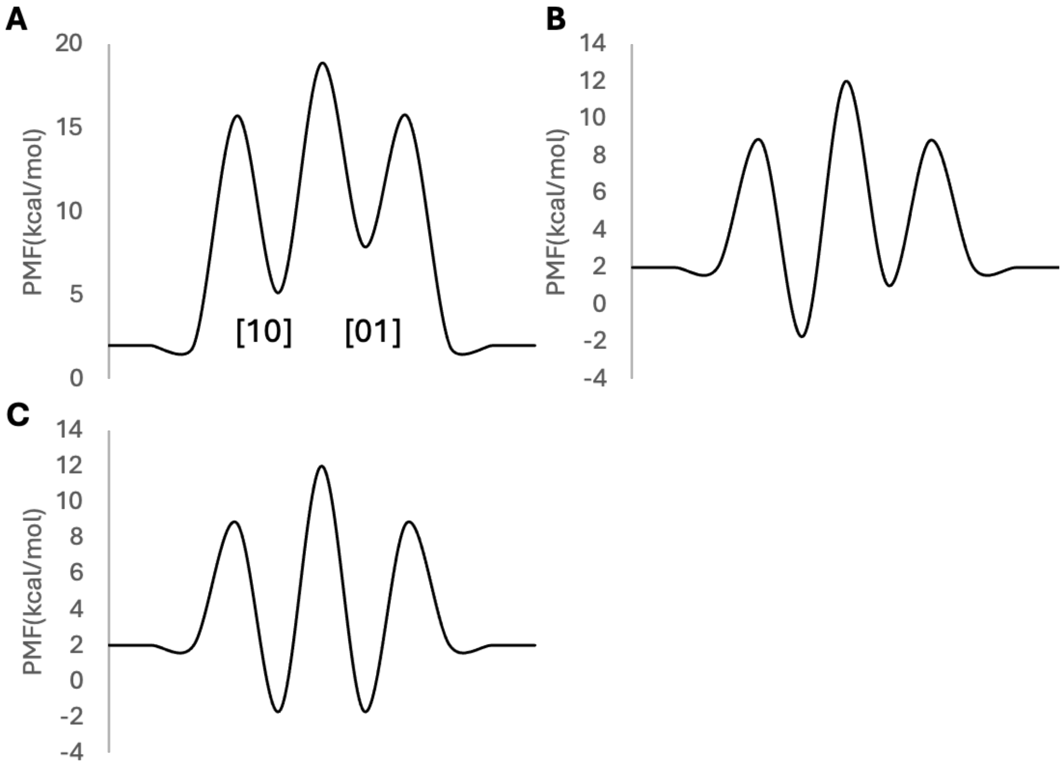


Figure S1.  Additional 2 site systems tested. A) Asymmetrical binding sites with wells less stable than the bulk solution. B) Asymmetrical binding sites with wells more stable than the bulk solution. C) Symmetrical binding sites with wells more stable than bulk solution.

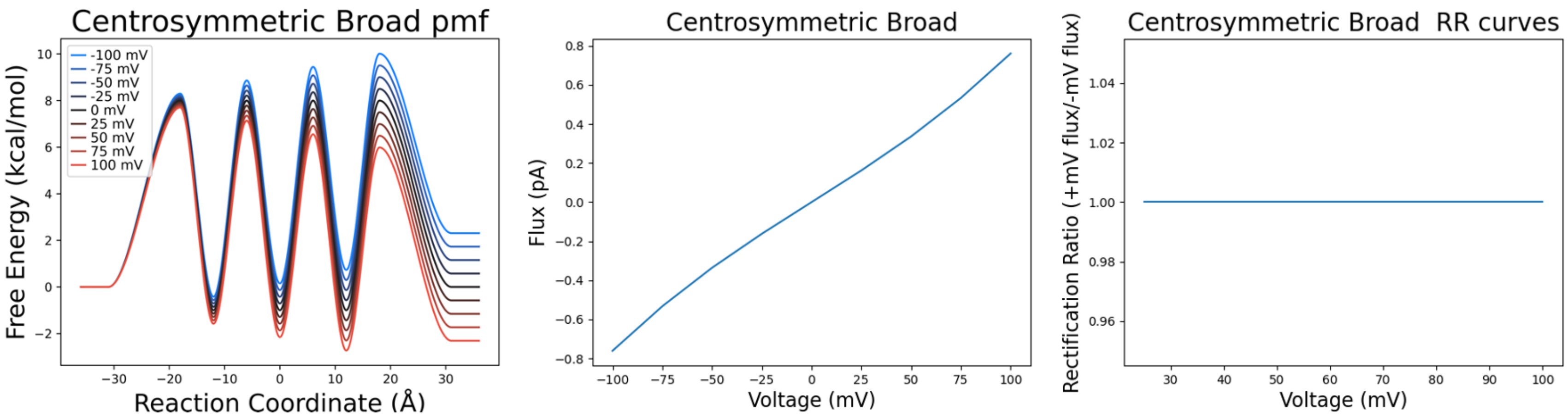


Figure S2. Symmetric distribution of site locations using a broader spread with binding sites at -12, 0, and 12 Å. Barriers located at -18, -6, 6, and 18 Å. No rectification was observed.

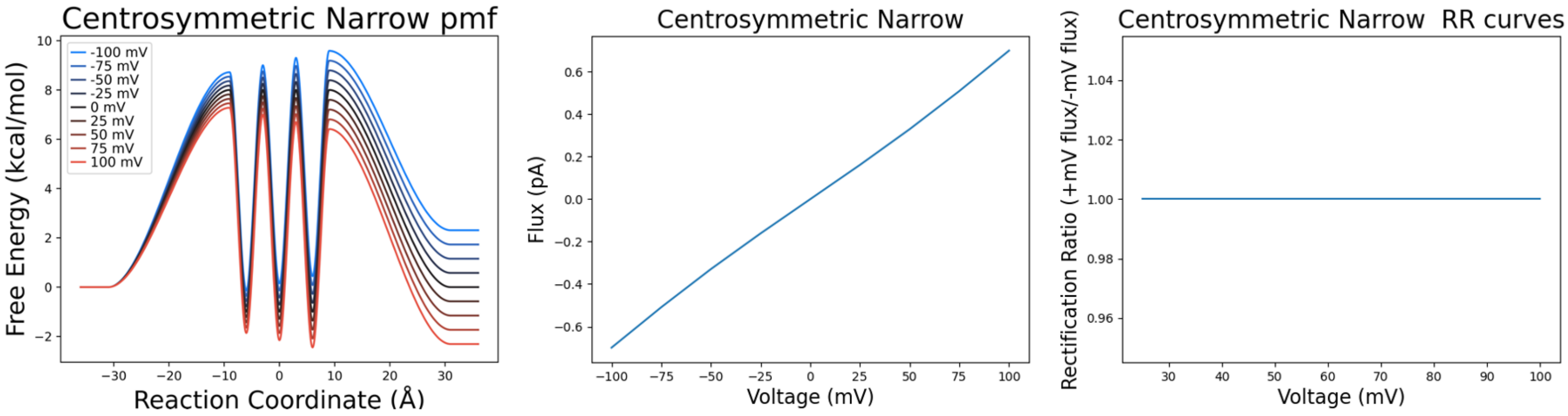


Figure S3. Symmetric distribution of site locations using a narrower spread with binding sites at -6, 0, and 6 Å. Barriers located at -9, -3, 3, and 9 Å. No rectification was observed.

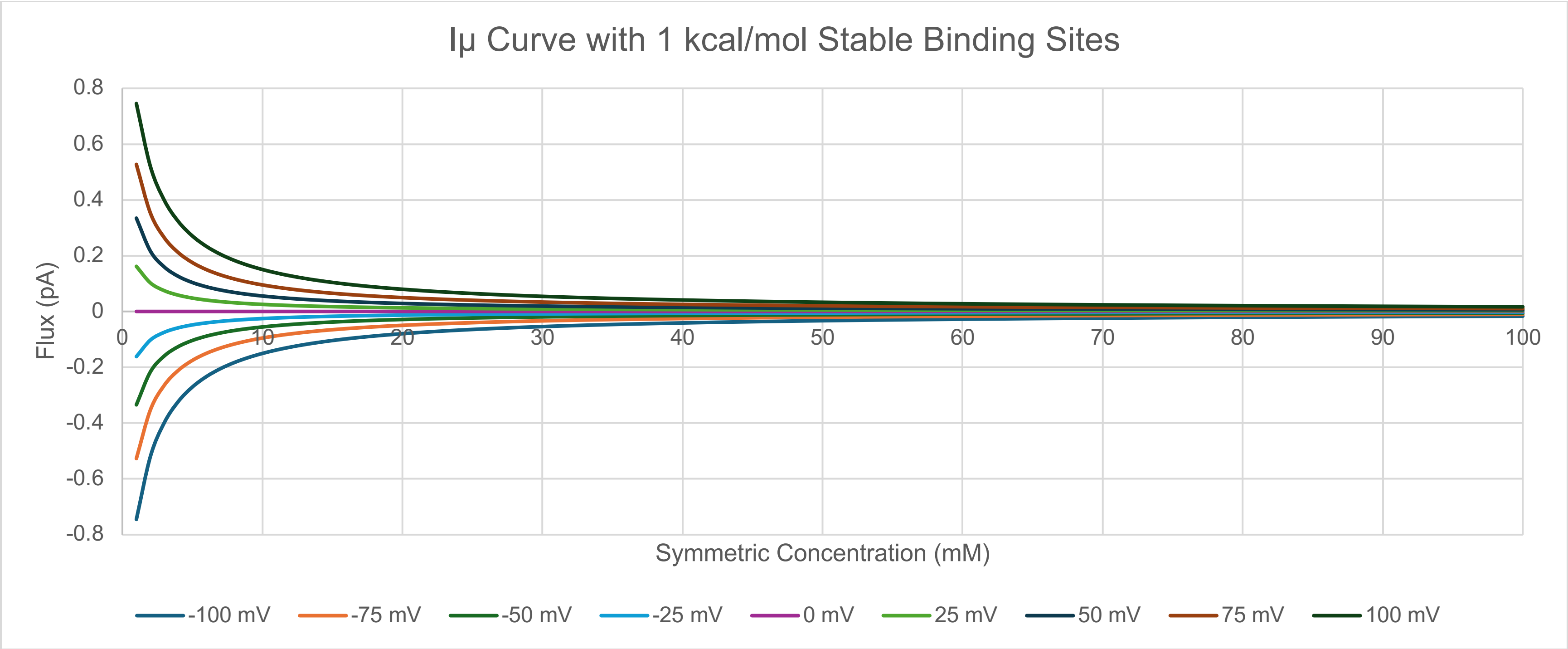

Figure S4. Iμ curve when binding sites are more stable than bulk shows immediate decrease in flux with increasing concentration, but quickly saturates.

Table S1. Symmetrical binding sites with wells less stable than the bulk solution (Fig. 5A).

| Driving Force(s) | -140mV | 199.15: 0.85 | -70mV | 158.7: 10.3 | 0mV 100:100 | +70mV | 10.3 :158.7 | +140mV | 0.85mM : 199.15mM |
|---|---|---|---|---|---|---|---|---|---|
| Net Flux | 2.60E+00 | 4.91E-01 | 8.21E-01 | 3.98E-01 | 0.00E+00 | -8.21E-01 | -3.98E-01 | -2.60E+00 | -4.91E-01 |
| Pop [00] | 3.57E-01 | 5.00E-01 | 4.57E-01 | 5.42E-01 | 5.00E-01 | 4.57E-01 | 5.42E-01 | 3.57E-01 | 5.00E-01 |
| Pop [01] | 5.84E-01 | 4.95E-01 | 4.16E-01 | 4.28E-01 | 2.50E-01 | 1.27E-01 | 2.99E-02 | 5.93E-02 | 4.58E-03 |
| Pop [10] | 5.93E-02 | 4.58E-03 | 1.27E-01 | 2.99E-02 | 2.50E-01 | 4.16E-01 | 4.28E-01 | 5.84E-01 | 4.95E-01 |

[int]:[ext] in mM

Table S2: Asymmetrical binding sites with wells less stable than the bulk solution (Fig. S1A).

| Driving Force(s) | -140mV | 199.15: 0.85 | -70mV | 158.7: 10.3 | 0mV 100:100 | +70mV | 10.3 :158.7 | +140mV | 0.85mM : 199.15mM |
|---|---|---|---|---|---|---|---|---|---|
| Net Flux | 6.17E+00 | 4.91E-01 | 1.40E+00 | 3.98E-01 | 0.00E+00 | -9.39E-01 | -3.98E-01 | -2.76E+00 | -4.91E-01 |
| Pop [00] | 8.45E-01 | 9.81E-01 | 7.77E-01 | 9.41E-01 | 6.64E-01 | 5.23E-01 | 5.59E-01 | 3.79E-01 | 5.02E-01 |
| Pop [01] | 1.38E-02 | 9.72E-03 | 7.08E-03 | 7.43E-03 | 3.32E-03 | 1.45E-03 | 3.08E-04 | 6.30E-04 | 4.60E-05 |
| Pop [10] | 1.41E-01 | 8.99E-03 | 2.16E-01 | 5.19E-02 | 3.32E-01 | 4.76E-01 | 4.41E-01 | 6.20E-01 | 4.98E-01 |

[int]:[ext] in mM

Table S3: Asymmetrical binding sites with wells more stable than the bulk solution (Fig. S1B).

| Driving Force(s) | -140mV | 199.15: 0.85 | -70mV | 158.7: 10.3 | 0mV 100:100 | +70mV | 10.3 :158.7 | +140mV | 0.85mM : 199.15mM |
|---|---|---|---|---|---|---|---|---|---|
| Net Flux | 3.99E+01 | 5.15E+01 | 6.27E+00 | 1.16E+01 | 0.00E+00 | -1.97E+00 | -9.29E-01 | -4.45E+00 | -9.91E-01 |
| Pop [00] | 5.47E-05 | 5.24E-04 | 3.49E-05 | 3.49E-05 | 1.98E-05 | 1.09E-05 | 1.27E-05 | 6.10E-06 | 1.01E-05 |
| Pop [01] | 8.96E-02 | 5.19E-01 | 3.18E-02 | 3.18E-02 | 9.90E-03 | 3.04E-03 | 6.98E-04 | 1.02E-03 | 9.24E-05 |
| Pop [10] | 9.10E-01 | 4.80E-01 | 9.68E-01 | 9.68E-01 | 9.90E-01 | 9.97E-01 | 9.99E-01 | 9.99E-01 | 1.00E+00 |

[int]:[ext] in mM

Table S4: Symmetrical binding sites with wells more stable than the bulk solution (Fig. S1C).

| Driving Force(s) | -140mV | 199.15: 0.85 | -70mV | 158.7: 10.3 | 0mV 100:100 | +70mV | 10.3 :158.7 | +140mV | 0.85mM : 199.15mM |
|---|---|---|---|---|---|---|---|---|---|
| Net Flux | 4.04E+00 | 9.82E-01 | 1.51E+00 | 8.69E-01 | 0.00E+00 | -1.51E+00 | -8.69E-01 | -4.04E+00 | -9.82E-01 |
| Pop [00] | 5.54E-06 | 1.00E-05 | 8.41E-06 | 1.18E-05 | 1.00E-05 | 8.41E-06 | 1.18E-05 | 5.54E-06 | 1.00E-05 |
| Pop [01] | 9.08E-01 | 9.91E-01 | 7.67E-01 | 9.35E-01 | 5.00E-01 | 2.33E-01 | 6.53E-02 | 9.22E-02 | 9.16E-03 |
| Pop [10] | 9.22E-02 | 9.16E-03 | 2.33E-01 | 6.53E-02 | 5.00E-01 | 7.67E-01 | 9.35E-01 | 9.08E-01 | 9.91E-01 |

[int]:[ext] in mM

Table S5:  Nernst cancelation for symmetrical binding sites with wells less stable than the bulk solution (Fig 5A).

| Driving Force(s) | -140mV + 0.85:199.15 | -70mV + 10.3:158.7 | +70mV + 158.7:10.3 | +140mV + 199.15:0.85 |
|---|---|---|---|---|
| Net Flux | -3.27E-05 | -3.99E-05 | 3.99E-05 | 3.27E-05 |
| Pop [00] | 7.66E-01 | 6.56E-01 | 6.56E-01 | 7.66E-01 |
| Pop [01] | 1.11E-02 | 6.25E-02 | 2.81E-01 | 2.23E-01 |
| Pop [10] | 2.23E-01 | 2.81E-01 | 6.25E-02 | 1.11E-02 |

[int]:[ext] in mM

Table S6: Nernst cancelation for asymmetrical binding sites with wells less stable than the bulk solution (Fig S1A).

| Driving Force(s) | -140mV + 0.85:199.15 | -70mV + 10.3:158.7 | +70mV + 158.7:10.3 | +140mV + 199.15:0.85 |
|---|---|---|---|---|
| Net Flux | -3.31E-05 | -4.25E-05 | 5.52E-05 | 4.20E-05 |
| Pop [00] | 7.74E-01 | 6.99E-01 | 9.09E-01 | 9.83E-01 |
| Pop [01] | 1.12E-04 | 6.66E-04 | 3.90E-03 | 2.87E-03 |
| Pop [10] | 2.26E-01 | 3.00E-01 | 8.67E-02 | 1.43E-02 |

[int]:[ext] in mM

Table S7: Nernst cancelation for asymmetrical binding sites with wells more stable than the bulk solution (Fig S1B).

| Driving Force(s) | -140mV + 0.85:199.15 | -70mV + 10.3:158.7 | +70mV + 158.7:10.3 | +140mV + 199.15:0.85 |
|---|---|---|---|---|

| | | | | |
|---|---|---|---|---|
| Net Flux | -1.47E-04 | -1.41E-04 | 6.10E-04 | 2.45E-03 |
| Pop [00] | 3.43E-05 | 2.33E-05 | 1.00E-04 | 5.74E-04 |
| Pop [01] | 4.97E-04 | 2.22E-03 | 4.31E-02 | 1.67E-01 |
| Pop [10] | 9.99E-01 | 9.98E-01 | 9.57E-01 | 8.32E-01 |

[int]:[ext] in mM

Table S8: Nernst cancelation for symmetrical binding sites with wells more stable than the bulk solution (Fig S1C).

| Driving Force(s) | -140mV + 0.85:199.15 | -70mV + 10.3:158.7 | +70mV + 158.7:10.3 | +140mV + 199.15:0.85 |
|---|---|---|---|---|
| Net Flux | -1.40E-04 | -1.16E-04 | 1.16E-04 | 1.40E-04 |
| Pop [00] | 3.27E-05 | 1.91E-05 | 1.91E-05 | 3.27E-05 |
| Pop [01] | 4.74E-02 | 1.82E-01 | 8.18E-01 | 9.53E-01 |
| Pop [10] | 9.53E-01 | 8.18E-01 | 1.82E-01 | 4.74E-02 |

[int]:[ext] in mM

Table S9: Symmetrical binding sites with wells more stable than the bulk solution and site 1 moved 10A towards center.

| Driving Force(s) | -140mV | 199.15: 0.85 | -70mV | 158.7: 10.3 | 0mV 100:100 | +70mV | 10.3 :158.7 | +140mV | 0.85mM : 199.15mM |
|---|---|---|---|---|---|---|---|---|---|
| Net Flux | 1.45E+00 | 9.82E-01 | 9.73E-01 | 8.69E-01 | 0.00E+00 | -1.67E+00 | -8.69E-01 | -4.29E+00 | -9.82E-01 |
| Pop [00] | 1.95E-06 | 1.00E-05 | 5.39E-06 | 1.18E-05 | 1.00E-05 | 9.32E-06 | 1.18E-05 | 5.89E-06 | 1.00E-05 |
| Pop [01] | 9.67E-01 | 9.91E-01 | 8.50E-01 | 9.35E-01 | 5.00E-01 | 1.51E-01 | 6.53E-02 | 3.58E-02 | 9.16E-03 |
| Pop [10] | 3.26E-02 | 9.16E-03 | 1.50E-01 | 6.53E-02 | 5.00E-01 | 8.49E-01 | 9.35E-01 | 9.64E-01 | 9.91E-01 |

[int]:[ext] in mM

Table S10: Nernst cancelation for symmetrical binding sites with wells more stable than the bulk solution and site 1 moved 10A towards center (right).

| Driving Force(s) | -140mV + 0.85:199.15 | -70mV + 10.3:158.7 | +70mV + 158.7:10.3 | +140mV + 199.15:0.85 |
|---|---|---|---|---|
| Net Flux | -1.30E-04 | -1.03E-04 | 1.77E-04 | 3.80E-04 |

| | | | | |
|---|---|---|---|---|
| Pop [00] | 2.99E-05 | 1.68E-05 | 2.91E-05 | 8.89E-05 |
| Pop [01] | 1.29E-01 | 2.77E-01 | 7.23E-01 | 8.71E-01 |
| Pop [10] | 8.71E-01 | 7.23E-01 | 2.77E-01 | 1.29E-01 |

[int]:[ext] in mM

Table S11: Symmetrical binding sites with wells more stable than the bulk solution and site 1 moved 10A towards bulk (left).

| Driving Force(s) | -140mV | 199.15: 0.85 | -70mV | 158.7: 10.3 | 0mV 100:100 | +70mV | 10.3 :158.7 | +140mV | 0.85mM : 199.15mM |
|---|---|---|---|---|---|---|---|---|---|
| Net Flux | 7.32E+00 | 9.82E-01 | 1.94E+00 | 8.69E-01 | 0.00E+00 | -1.38E+00 | -8.69E-01 | -3.73E+00 | -9.82E-01 |
| Pop [00] | 1.02E-05 | 1.00E-05 | 1.08E-05 | 1.18E-05 | 1.00E-05 | 7.69E-06 | 1.18E-05 | 5.11E-06 | 1.00E-05 |
| Pop [01] | 8.31E-01 | 9.91E-01 | 7.00E-01 | 9.35E-01 | 5.00E-01 | 2.99E-01 | 6.53E-02 | 1.62E-01 | 9.16E-03 |
| Pop [10] | 1.69E-01 | 9.16E-03 | 3.00E-01 | 6.53E-02 | 5.00E-01 | 7.01E-01 | 9.35E-01 | 8.38E-01 | 9.91E-01 |

[int]:[ext] in mM

Table S12: Nernst cancelation symmetrical binding sites with wells more stable than the bulk solution and site 1 moved 10A towards bulk (left).

| Driving Force(s) | -140mV + 0.85:199.15 | -70mV + 10.3:158.7 | +70mV + 158.7:10.3 | +140mV + 199.15:0.85 |
|---|---|---|---|---|
| Net Flux | -1.41E-04 | -1.22E-04 | 8.69E-05 | 7.23E-05 |
| Pop [00] | 3.34E-05 | 2.01E-05 | 1.43E-05 | 1.69E-05 |
| Pop [01] | 2.45E-02 | 1.36E-01 | 8.64E-01 | 9.75E-01 |
| Pop [10] | 9.75E-01 | 8.64E-01 | 1.36E-01 | 2.45E-02 |

[int]:[ext] in mM
